# A High-Resolution Luminescent Assay for Rapid and Continuous Monitoring of Protein Translocation Across Biological Membranes

**DOI:** 10.1101/456921

**Authors:** Gonçalo C. Pereira, William J. Allen, Daniel W. Watkins, Lisa Buddrus, Dylan Noone, Xia Liu, Andrew Richardson, Agnieszka Chacinska, Ian Collinson

## Abstract

Protein translocation is a fundamental process in biology. Major gaps in our understanding of this process arises due the poor sensitivity, low time-resolution and irreproducibility of translocation assays. To address this, we applied NanoLuc split-luciferase to produce a new strategy for measuring protein transport. The system reduces the timescale of data collection from days to minutes, and allows continuous acquisition with a time-resolution in the order of seconds – yielding kinetics parameters suitable for mechanistic elucidation and mathematical fitting. To demonstrate its versatility, we implemented and validated the assay *in vitro* and *in vivo* for the bacterial Sec system, and the mitochondrial protein import apparatus. Overall, this technology represents a major step forward, providing a powerful new tool for fundamental mechanistic enquiry of protein translocation and for inhibitor (drug) screening, with an intensity and rigour unattainable through classical methods.

## INTRODUCTION

A large proportion of proteins fulfil their function outside the cell, or in a subcellular compartment distinct from the cytosol. To get there, they must be sorted and then transported across the appropriate membranes; thus, protein translocation systems are ubiquitous and fundamental features of cellular compartmentalisation. For example, bacteria target about 20% of their proteome to the Sec system for transport across or into the plasma membrane^1^, while nearly all mitochondrial proteins are produced in the cytosol for import through the **T**ranslocases of the **O**uter and **I**nner **M**embranes (TOM and TIM). In both of these cases, proteins are recognised by specific targeting sequences that are often cleaved upon completion of transport to liberate the mature protein.

Previously, protein translocation has been monitored *in vitro* by quantifying time courses of proteins transported into the interior of reconstituted proteoliposomes (PLs) or native membranes, *e.g.* bacterial inner membrane vesicles^2^^,^^3^ (IMVs) or intact mitochondria^4^^,^^5^ In these experiments, successfully transported protein is typically characterised by resistance to proteolysis and detected by Western blotting or autoradiography. These classical methods have been instrumental for the determination of the molecular components and basic properties of the various prokaryotic^6^ and eukaryotic translocation apparatus^7^^,^^8^. However, such assays are not suited to a more sophisticated analysis, due to lack of kinetic detail – they produce only discontinuous, endpoint measurements – and are labour intense, making them difficult to scale to high-throughput. Over the past two decades, various alternative methods have been proposed (Table S1), all of which have drawbacks. A highly sensitive, versatile and quantitative real-time assay has yet to be developed.

Recently, Dixon, *et al.* ^9^ developed a non-covalent complementation system based on a small, bright luciferase, to monitor protein-protein interactions – NanoBiT (short for NanoLuc Binary Technology). In this split-luciferase, the final β-strand of NanoLuc was cleaved to generate a large fragment of 18 kDa, referred to as *11S* (trademark name *LgBiT*), and a small 1.3 kDa peptide chain of 11 amino acids, termed *pep114* (trademark name *SmBiT*). The authors also developed a high-affinity variant of pep114, *pep86* (trademark name *HiBiT*; picomolar range); here, we exploit the rapid, spontaneous interaction between 11S and pep86 as the basis for a protein translocation assay.

Using simple genetic tools, we targeted 11S to two model destination compartments: the yeast mitochondrial matrix and the *Escherichia coli* periplasm (Fig. 1). The presence of lipid bilayers keeps the reporter segregated, ensuring that complementation is restricted to the destination cell compartment. Substrates pre-proteins, *i.e.* proteins with their targeting sequence uncleaved, are then tagged with pep86 at their C-terminus. Upon translocation, the rise in local protein concentration leads to complementation of pep86 with internalised 11S, producing active luciferase activity and thus a readout for protein translocation (Fig. 1). Pep86 is small and native-like, so should eliminate artefacts caused by non-native tags such as fluorescent dyes, and have negligible effect on transport rates. The enzymatic amplification generated by NanoLuc, meanwhile, provides very high sensitivity measurements compared to conventional methods (Table S1), allowing detailed quantitative analyses of translocation. We anticipate that the techniques and tools described here will readily be transferrable to many other membrane and non-membrane protein translocation systems.

**Figure 1.**
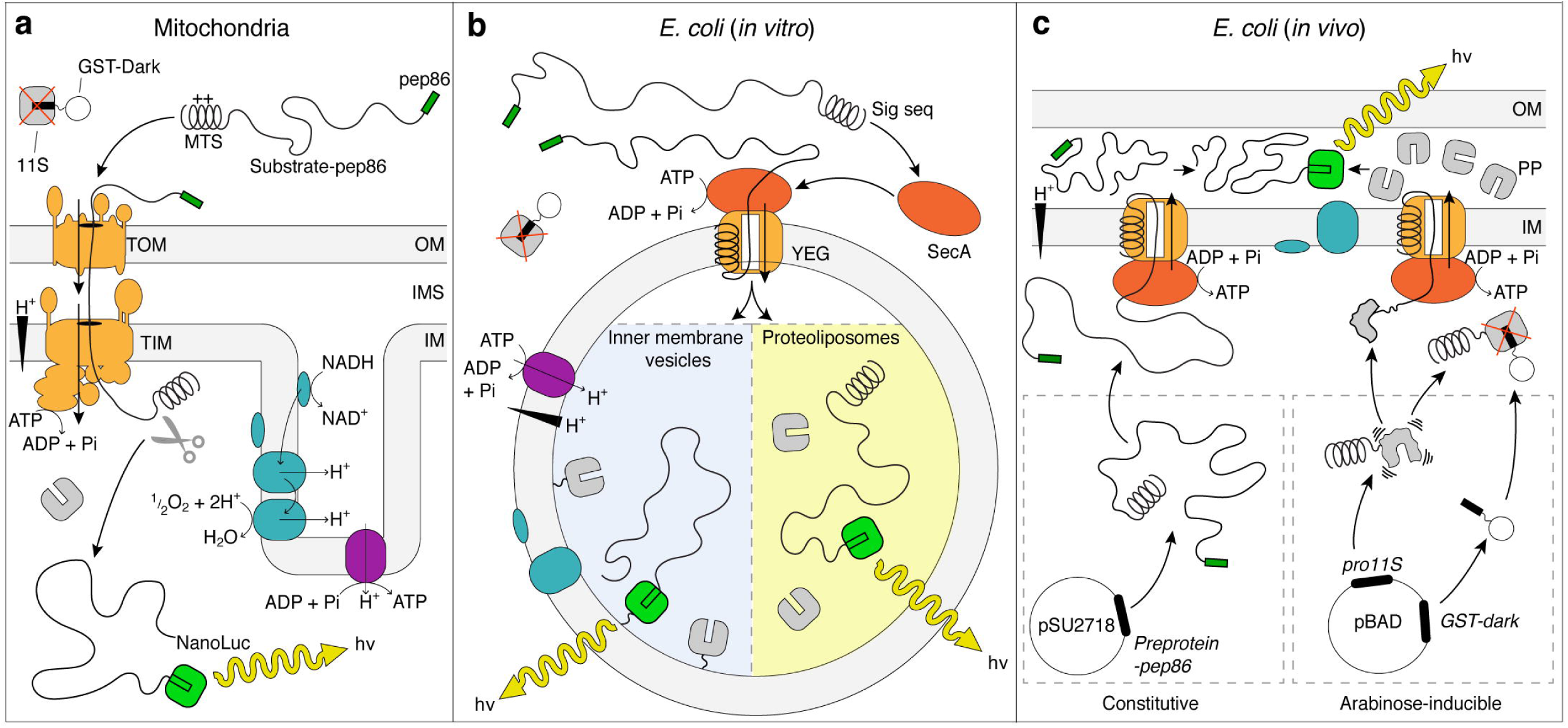
Overview of the new real-time assay to monitor protein translocation on different biological systems. The large 11S fragment was segregated in the mitochondrial matrix, proteoliposome lumen, tethered to the inner membrane of *E. coli* (inner membrane vesicles) or in the periplasm (*in vivo*). All pre-protein substrates were tagged on their C-terminus with the high-affinity peptide pep86. To decrease signal background, a non-luminescent high-affinity pep86 peptide (GST-dark) was added to the reactions (mitochondria, *E. coli in vitro*) or co-expressed in the cytosol for *in vivo* experiments. In all systems, successful pre-protein translocation was observed as an increase in luminescent signal upon pep86—11S complementation. Abbreviations: *hv* – light; IM – inner membrane; IMS – intermembrane sequence; MTS – mitochondrial targeting sequence; OM – outer membrane; PP – periplasm; Sig. seq. – signal sequence; TIM – translocase of inner membrane; TOM – translocase of outer membrane.

## RESULTS

### *In vitro* continuous translocation assay of the bacterial Sec machinery

The bacterial Sec system is the principal mechanism for protein secretion across the bacterial plasma membrane, and a good starting point for assay development as its activity has been extensively studied. Transport through the Sec machinery can be recapitulated *in vitro* using only a small number of purified components: PLs^6^ or IMVs^10^^,^^11^ containing the SecYEG protein-channel membrane protein complex, the soluble motor ATPase SecA, a pre-secretory protein substrate with a cleavable N-terminal signal sequence and ATP as an energy source. In both cases, successful translocation results in internalisation of pre-protein, equivalent to translocation into the periplasm *in vivo*. When the reaction is complete, protease K is added to digest untranslocated pre-protein, while the vesicle protects the successfully translocated material. Alternatively, samples can be taken from the reaction at various time points and quenched with ice-cold buffer containing protease K, to investigate the transport kinetics. Analysis is typically performed by SDS-PAGE followed by autoradiography (of radiolabelled substrates) or immunoblotting (Fig. 2a).

**Figure 2.**
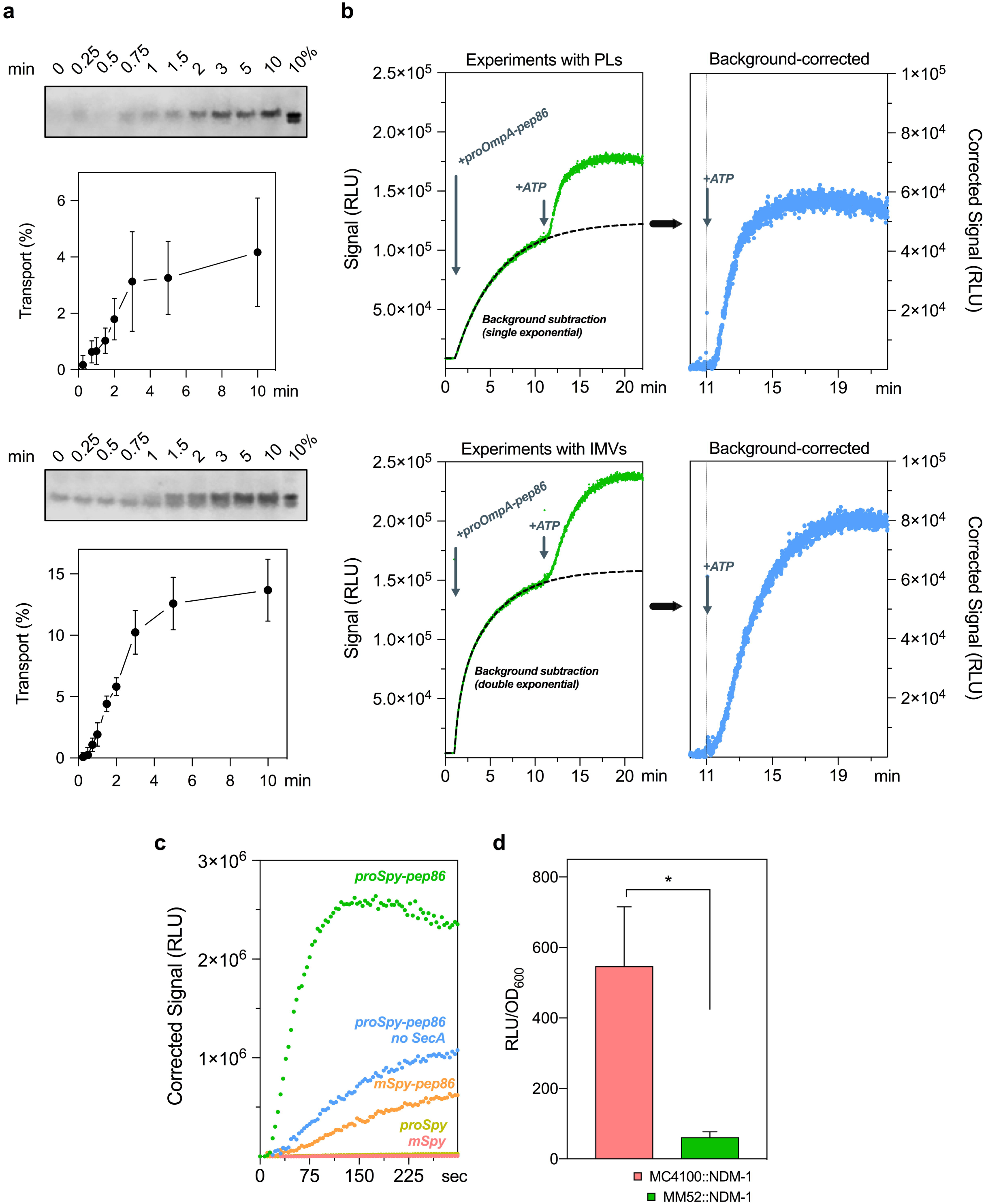
Comparison of conventional *vs*. real-time translocation assay for the bacterial SecYEG sytem. **A** – in vitro reactions in PLs (top) and IMVs (bottom) were carried in the presence of an ATP regenerating system, SecA and 200 nM of proOmpA; reaction started by addition of 1mM ATP. At the indicated timepoints, an aliquot was withdrawn and quickly quenched by dilution in ice-cold protease K and EDTA. Protease-protected OmpA was evaluated by SDS-PAGE followed by immunoblotting. **B –** for the real-time translocation assay, 11S (40 µM) was included in the buffers for PLs preparation ensuring its encapsulation (top) or tethered to the inner membrane of HB1 cells (IMVs; bottom). Reactions were carried out on a cuvette luminometer with identical buffers as described before. Reactions were started by addition of proOmpA in the absence of ATP and the luminescence background was monitored, then ATP was added to start pre-protein translocation. **C** - Real-time import of proSpy-pep86 into BL21(DE3) IMVs containing 11S. Reactions were carried out as described in (A) but on 96 well-plate format. Plot represents the signal after background is omitted from the graph. Deletion of the signal sequence (mSpy-pep86) or absence of SecA prevented import of pre-protein. No signal is observed if pep86 is absent from the pre-protein. **D** – evaluation of NDM-1-pep86 translocation *in vivo* by co-expressing periplasm-targeted 11S; end-point measurements were taken after 2 h induction and compared to a SecA-temperature sensitive mutant (MM52 strain) and normalised to cell number. Abbreviations: IMVs – inner membrane vesicles; PLs – proteoliposomes.

These assays do not enable an exploration beyond a simple analysis, so we set out to modify SecYEG PLs, IMVs and substrate pre-proteins such that they would be compatible with a split NanoLuc-based assay (Fig. 1b). For PLs, we included purified _6H_11S into the reconstitution mixture then removed excess protein by three rounds of centrifugation and resuspension in clean buffer. To ensure a high concentration of internalised 11S within IMVs, we produced a variant of 11S with an inner membrane lipid anchor sequence^12^. We then co-transformed the expression vectors harbouring 11S and SecYEG into *E. coli* and induced expression of both plasmids simultaneously. Pre-proteins were extended at the C-terminus by a GSG linker followed by pep86.

Reactions were initially set up with either PLs or IMVs saturated with SecA together with: an ATP regeneration system, Prionex – a biocompatible polymer used for protein stabilisation and to prevent 11S/pre-protein-pep86 from adhering to surfaces, and the NanoLuc substrate furimazine. Luminescence measurements were then started to generate a baseline reading, followed by addition of the model pre-protein, proOmpA, fused to pep86 (Figure 2b).

The initial experiments were prone to a high, ATP-independent luminescence signal, presumably due to contaminating external 11S, remnants of the reconstitution or from burst vesicles. This problem was resolved by supplementing the mixture with an inactivated form of the pep86, ‘*dark*’ peptide^*13*^. This peptide differs from the pep86 sequence by a point mutation at a critical catalytic Arginine residue, which does not prevent high affinity binding to 11S, but prevents catalysis once bound. For ease of production, we synthesised a fusion of glutathione-S-transferase and the ‘*dark*’ peptide, which we named *GST-dark*. The inclusion of 40 µM GST-dark did indeed massively reduce, but did not completely obviate, the background signal contribution (Fig. 2b). The remaining background signal could be fitted to either a single or double exponential function and subtracted from the data acquired after the addition of ATP (Fig. 2b).

The high detail of the background-corrected transport traces is clear (Fig. 2b). To confirm that the data report on the kinetics of translocation rather than slow rate-limiting pep86-11S association, we performed further assays with PLs containing a range of 11S concentrations (Fig. S1). The results show that while the signal amplitude is proportional to the concentration of 11S – *i. e.* the reaction ends when 11S runs out – the shape of the curve is completely unaffected. Thus, rates extracted from the data do indeed reflect the rate of transport.

To demonstrate the versatility of the assay, we also linked the pep86 sequence to the C-terminus of a very different *E. coli* pre-protein – the soluble, positively charged and α-helical spheroplast protein Y (Spy), for comparison against the β-barrelled proOmpA. The translocation data for proSpy (Fig. 2c, green trace) are qualitatively similar to those of proOmpA, demonstrating the broad compatibility of the assay. Importantly, removal of either the signal sequence or pep86 from proSpy, or SecA from the reaction mixture collapsed the signal. The omission of SecA from experiments conducted with IMVs retained residual activity (Fig 2c, blue trace), probably due to contamination by endogenous membrane associated SecA^14^.

Taken together, the results show that the luminescence signal is a *bona fide* measure of protein transport and suitable for a comprehensive kinetic analysis of the ATP and proton motive force (PMF) driven secretion process – to be described in forthcoming publications.

### *In vivo* β-lactamase secretion assay

Next, we set out to design a split-NanoLuc-based system for measuring translocation through the bacterial translocon *in vivo* (Fig. 1c). For this, we attached pep86 to the pre-secretory protein **N**ew **D**elhi **M**etallo-beta-lactamase 1 (NDM1) – a protein of great current interest due to its involvement in mediating antibiotic resistance in hospitals^15^. We also constructed an 11S variant with the N-terminal signal sequence of proOmpA (forming pro-11S), directing it to the periplasm. To reduce the background from newly synthesised pro-11S that has yet to be secreted, *GST-dark* was expressed in the cytoplasm (Fig. S2). By comparing an *E. coli* strain with severe secretion defects^16^ to its parent strain, we show that this system does indeed produce a secretion-dependent luminescence signal (Fig. 2d). This *in vivo* measure of secretion will be a powerful tool for understanding the native secretion process as well as for the development of new strategies for antibiotic discovery and against **A**nti-**M**icrobial **R**esistance (AMR).

### Real-time import assay in isolated yeast mitochondria

Over the past forty years, isolated mitochondrial fractions of *S. cerevisiae* have been used widely to study protein translocation *in vitro*. Since tools for genetic engineering of yeast are widespread and large quantities of the organism can be grown to yield considerable amounts of mitochondria, yeast mitochondria are a perfect candidate for applying split-NanoLuc to a eukaryotic protein translocation system (Fig. 1a).

Classical mitochondrial import assays employ autoradiography or western blotting (Fig. 3a) to detect transport. Similar to the bacterial setup, reaction aliquots representing time points were quenched by moving samples to ice followed by addition of a ‘death’ cocktail of oligomycin, valinomycin and antimycin A (OVA) and protease K, to respectively collapse the PMF and digest non-translocated precursors. Import is indicated by detection of the mature protein, *i.e.* after cleavage of the mitochondrial targeting sequence (MTS), by SDSPAGE followed by immunoblotting. As shown in Fig. 3a, the data points are discrete with a time resolution on the order of minutes.

**Figure 3.**
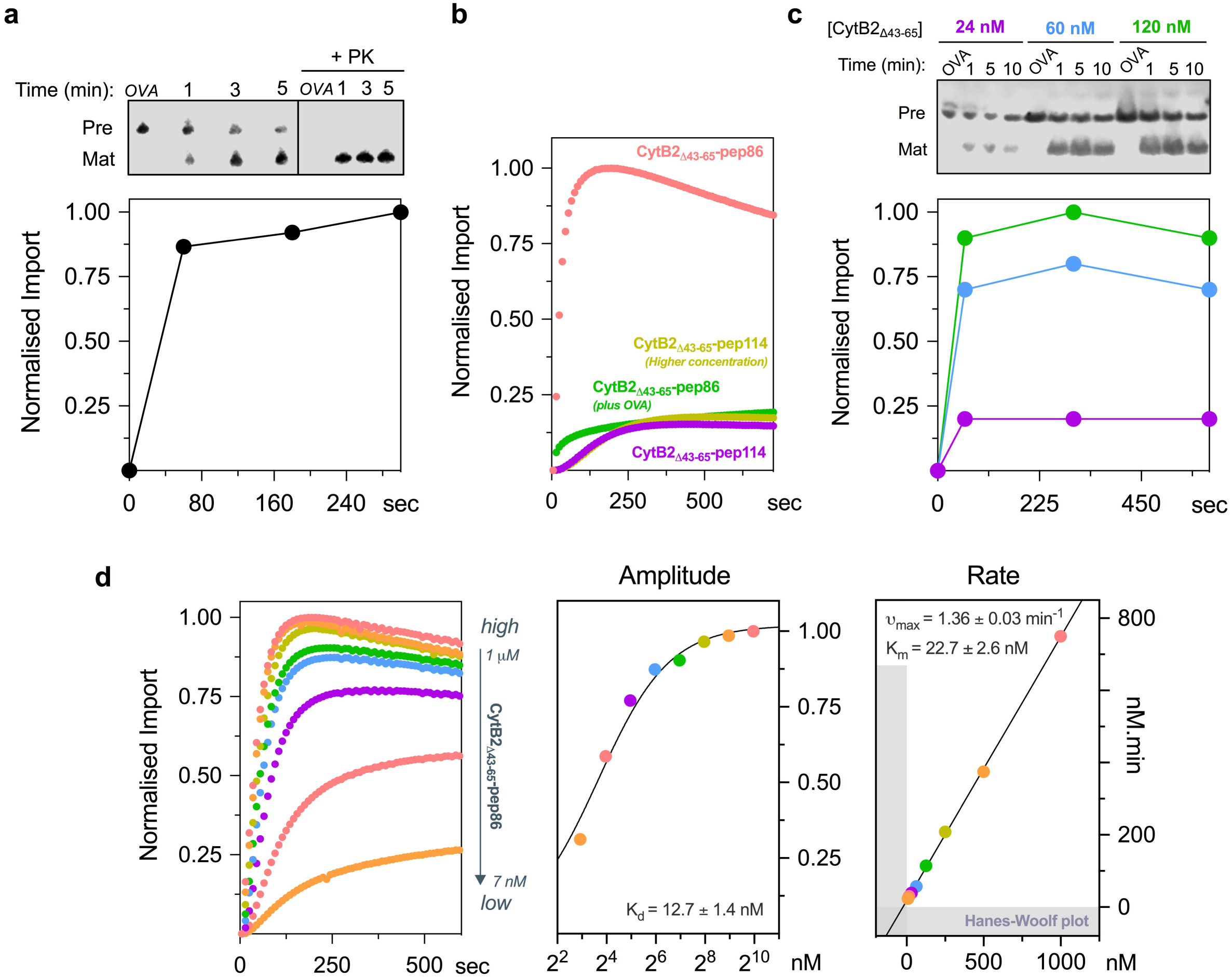
Comparison between the conventional method and the new real-time import assay to monitor protein import in mitochondria. **A**) conventional method -- fully energised mitochondria were challenged with 62 nM CytB2_Δ43-65_-pep86 and aliquots collected and quenched at specific timepoints. To identify successful translocation, samples were digested with protease K and mature (cleaved signal sequence) protein was identified by immunoblotting after SDS-PAGE. **B**) typical real-time import traces where fully energised mt-11S mitochondria were challenged with 1 µM CytB2_Δ43-65_-pep86 in the presence or absence of OVA. If the low affinity pep114 tag was used instead, no significant gain-of-signal was observed. (1 µM – low; 10 µM – high). Typical time-course of dose-response of CytB2_Δ43-65_-pep86 using the conventional (**C**) *vs.* the new real-time assay (**D**). The right panels on **D** represent parameters obtained from the normalised traces. The rate (*k*_app_) was calculated as the inverse of the time it takes to reach half of the maximal signal (t_50%_ in min). Abbreviations: OVA – oligomycin, valinomycin and antimycin cocktail.

To develop the new transport assay, we targeted 11S to the mitochondrial matrix using the presequence of subunit F_1_α of the yeast ATP synthase (mt-11S). After mitochondrial isolation, samples were analysed by SDS-PAGE to confirm localisation of 11S to the matrix (Fig. S3a). The mass of mt-11S is identical to that of purified _H6_11S, suggesting efficient localisation to the matrix, where the MTS is removed. Under standard culture conditions, 1% galactose resulted in 1.34 ± 0.05 µM of mature 11S in mitochondria, assuming a standard matrix volume of 1.1 µL/mg of protein^17^ and using pure _H6_11S as a standard (Fig. S3c).

As an import substrate, we used the classical yeast import precursor cytochrome B2 (CytB2; YML054C) with its hydrophobic sorting domain removed (CytB2_Δ43-65_), causing it to localise to the matrix^18^. We found that standard *in vitro* import reactions conditions^4^^,^^8^^,^^19^ required optimisation: but by lowering both the concentration of mitochondria and of furimazine, we were able to produce a strong, transport-dependent luminescence signal, which could be maintained for minutes (Fig. S4).

A typical optimised import assay trace is shown in Fig. 3b. It comprises a baseline corresponding to the background produced by 11S alone (omitted from the graph) followed by a sigmoidal shape upon CytB2_Δ43-65_-pep86 addition. No increase in signal was observed if the low-affinity tag pep114 was used, confirming that spontaneous complementation is necessary to report translocation (Fig. 3b, purple trace). In a separate experiment, mitochondrial respiration was inhibited, and membrane potential (ΔΨ) dissipated by addition of the OVA ‘death’ cocktail (Fig. 3b, green trace). Under these conditions, no rise in luminescence was observed upon addition of CytB2_Δ43-65_-pep86, confirming that the signal reflects energy-dependent import. Note that background luminescence in the presence of OVA was significantly higher than the baseline (~12%) – albeit without the characteristic sigmoidal shape (Fig.S5) – suggesting import-independent complementation is due to leakage of 11S from broken mitochondria.

To eliminate the possibility of mitochondrial poisons non-specifically interfering with the assay, we performed binding assays in the presence of low and high concentration of these drugs (Table S1). This was the case only for CCCP, which at 1 µM lowered the maximum amplitude to 40% if pre-incubated. Therefore, CCCP was not used from here onwards.

Next, we utilised the high-throughput character of the assay and monitored CytB2_Δ43-65_-pep86 import over a range of concentrations. Attempts to carry out a similar titration assay using the conventional method have failed to deliver reliable results (Fig. 3c), due to the limited range of concentrations that can be practically measured (~24-120 nM). By contrast, the split-luciferase assay produces a dose-response curves of CytB2_Δ43-65_-pep86 spanning two orders of magnitude (7-1000 nM), with sufficient data quality to extract kinetic parameters (Fig. 3d). To ensure the observed kinetics report on import, rather than complementation of the reporter, we performed binding experiments with CytB2_Δ43-65_-pep86 and _6H_11S alone. Although linking the pep86 tag to a precursor generally decreased its affinity for 11S, the K_d_ remains sufficiently low (33.5 ± 5.7 nM; Fig. S6) compared to the effective mt-11S concentration (1.34 ± 0.05 µM). Given that association of CytB2_Δ43-65_-pep86 and 11S is much faster than the rate of translocation (Fig. S6), we can be assured the signal provides a faithful measure of import.

The amplitude of import extracted from the dose-response curve of CytB2_Δ43-65_-pep86 was fitted to an equation for specific binding, yielding a low dissociation constant (K_d_) of 12.7 ± 1.4 nM. Furthermore, the import rates vary in a hyperbolic manner with respect to precursor concentration, suggesting a Michaelis-Menten relationship, shown in the linearized Hanes-Wolf plot (Fig. 3d). This confirms that CytB2_Δ43-65_-pep86 import is limited by the number of import sites.

### Exploring energy-dependency of the mitochondrial import system

Next, we explored the energy dependency of the mitochondrial import system, given that protein import is known to be ATP- and PMF-dependent. When mitochondria were energised with NADH in the absence of ATP, the addition of the mitochondrial respiration inhibitor antimycin A (AA) was able to inhibit CytB2_Δ43-65_-pep86 import, regardless of the concentration used (Fig. 4a). This demonstrates the requirement of a functional respiratory chain to generate PMF and drive import. Under this condition, de-energised mitochondria can hydrolyse ATP in an attempt to generate ΔΨ. Therefore, addition of ATP as part of an ATP regenerating system was able to restore import, suggesting that the reverse activity of ATP synthase generates sufficient ΔΨ to drive CytB2_Δ43-65_-pep86 import (Fig. 4b). Interestingly, higher concentrations of AA in this setup caused additional inhibition of import (amplitude and rate), which we attributed to non-specific effects on mitochondrial physiology. Further evidence for the ATP synthase driving import in the presence of AA was obtained when matrix ATP influx was blocked by the adenine nucleotide translocase inhibitor carboxyatractyloside (CAT; Fig. 4c) or ATP hydrolysis by the ATP synthase inhibited with oligomycin (Oligo; Fig. 4d). In the first case, ATP uptake is blocked, and the effect is two-fold – no ATP is available to drive import or to generate PMF. In the latter case, ATP enters the matrix but cannot be used to generate PMF due to inhibition by oligomycin. Although both drugs (CAT and Oligo) lowered the amplitude of import in a dose-dependent manner, oligomycin had no effect on k_app_ suggesting that ΔΨ determines the extent of CytB2_Δ43-65_ accumulation.

**Figure 4.**
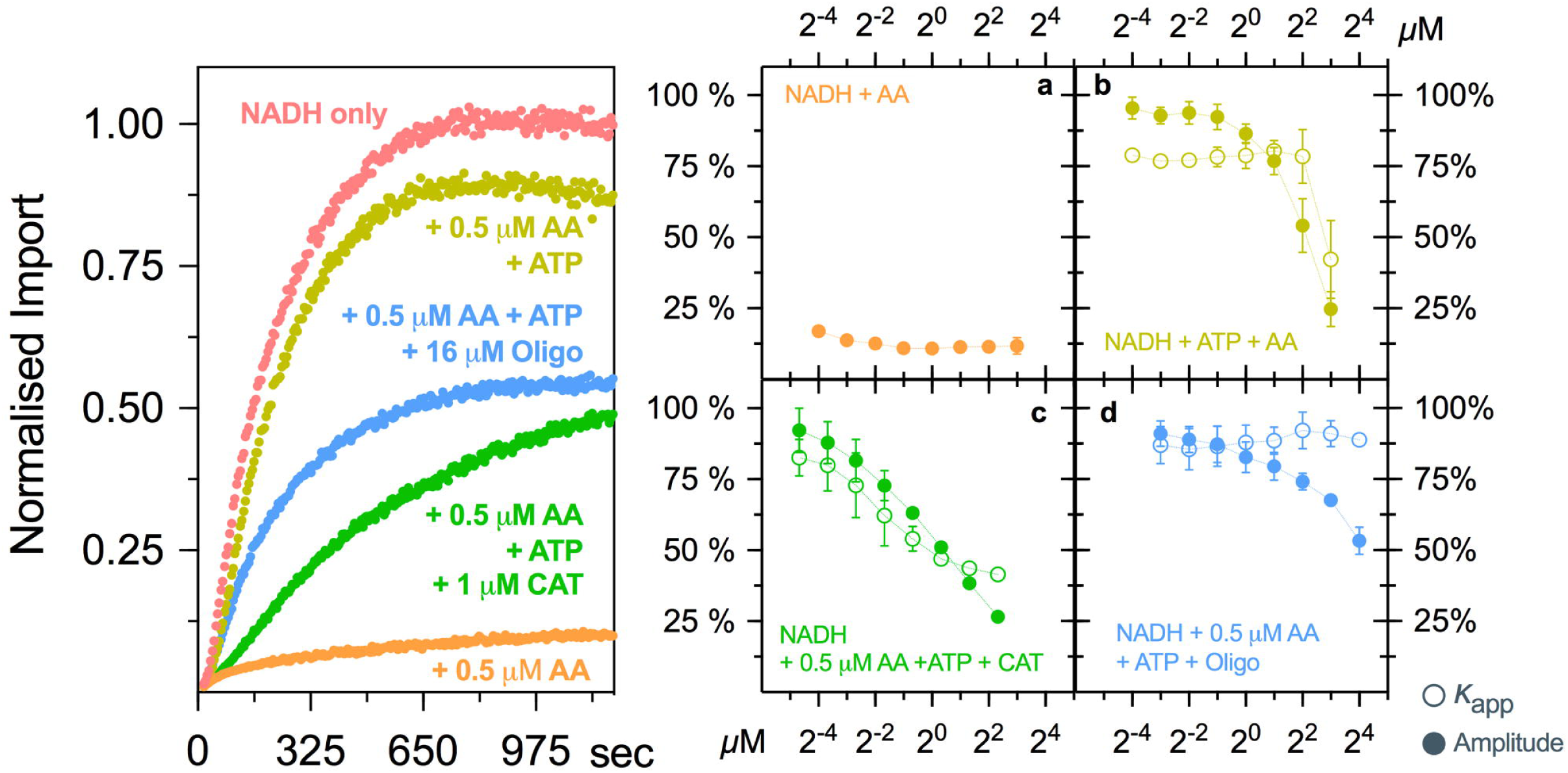
Energy-dependency of the mitochondrial import system. Mitochondria were energised in the presence of different mitochondrial poisons: antimycin A (AA; respiration inhibitior), oligomycin (Oligo; ATP synthase inhibitor) and/or carboxyatractyloside (CAT; adenine nucleotide translocase). Traces on the left, represent typical import reactions in the presence of these drugs. Dose-response curves are presented in the graphs on the right – full circles represent amplitude of import; empty circles show *k*_app._ Data is shown as mean ± SEM of 2-3 independent experiments. Error bars were omitted if smaller than symbols Abbreviations: AA – antimycin A; CAT – carboxyatractyloside; Oligo – oligomycin.

### Exploring the effect of small molecules on mitochondrial import

We used two small molecule inhibitors of the TIM23 pathway, identified by the Koehler Lab, to validate the assay: MB-12, also known as dequalinium^4^, and MB-10^5^. In accordance to previous reports^4^, we found that MB-12 causes a dose-dependent inhibition of CytB2_Δ43-65_-pep86 import with an IC_50_ 5.05 ± 0.32 µM (Fig. 5). The IC_50_ of MB-10, meanwhile, was 373 ± 16.4 µM (Fig. 5) – significantly higher than previous reported^5^. Interestingly, both drugs showed a stronger inhibitory effect on translocation when the mitochondria were energised by the reverse activity of the ATP synthase, possibly reflecting non-specific effects. This effect was more pronounced for MB-12 (IC_50_ = 0.437 ± 0.019 µM) than for MB-10 (IC_50_ = 102 ± 6.9 µM).

**Figure 5.**
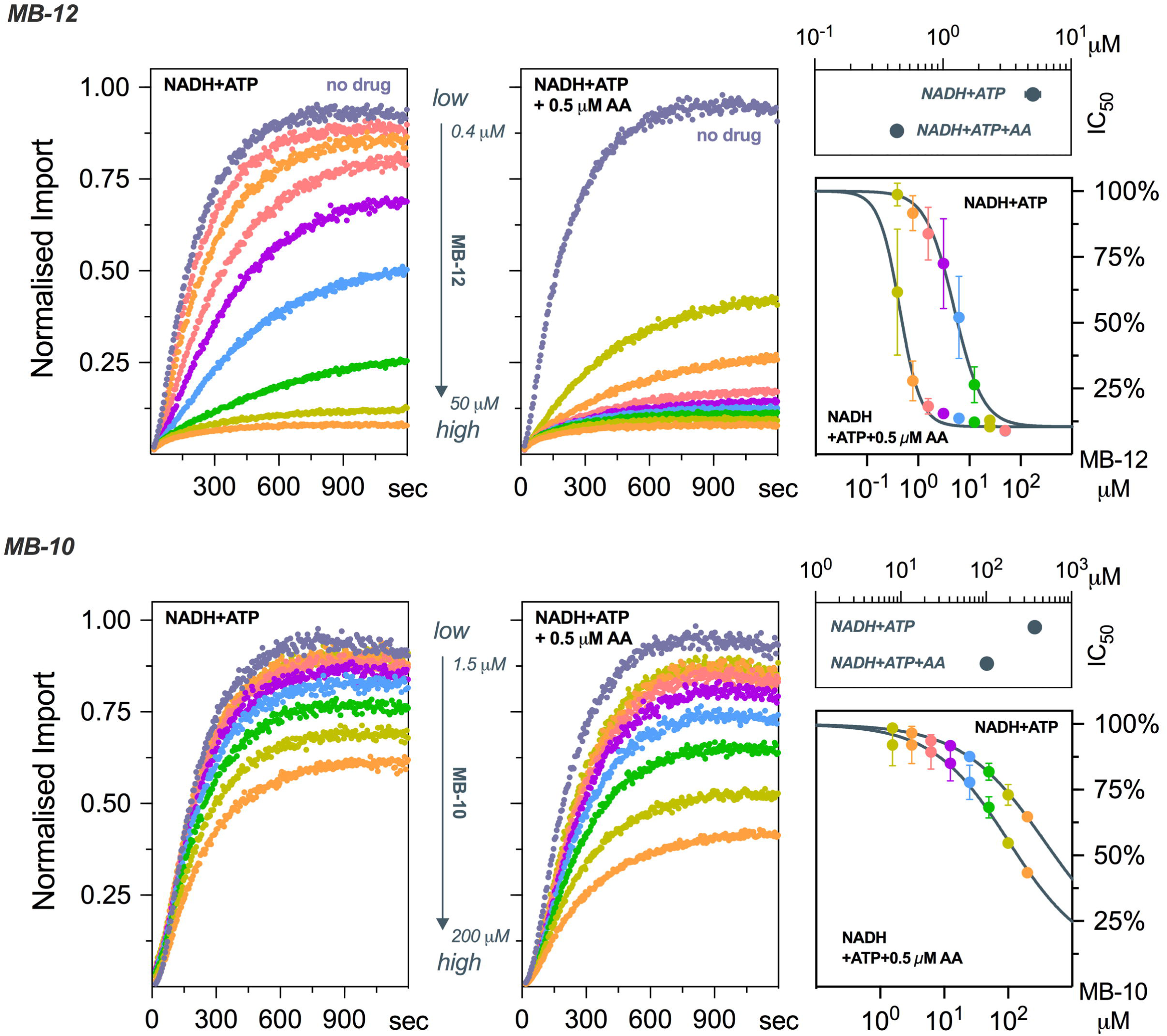
Effects of known-inhibitors of mitochondrial import. Dose-response curves of MB-12 (top) or MB-10 (bottom) on the import of CytB2_Δ43-65_-pep86 into mitochondria which PMF is created by NADH oxidation (left) or by the hydrolysis of ATP (right). Data is shown as mean ± SEM of three (MB-12) or two (MB-10) independent experiments, except for the IC_50_ where error bars represent 95% confidence intervals.

### Effect of signal sequence deletion

Finally, we evaluated the effect of removing the presequences from the import substrates, anticipating that the mature part alone would not import. However, because of the high sensitivity of our assay, we were able to observe that removal of the presequence^20^ (CytB2_Δ2-20 Δ43-65_-pep86) compromises but does not totally abolish import, as CytB2_Δ2-65_- or CytB2_Δ2-80_-pep86 do (Fig. 6). This residual activity reveals a potentially interesting new feature of the precursor targeting determinants: one that, along with many other facets of the import process, can now be dissected in detail by the application of this validated new and powerful tool.

**Figure 6.**
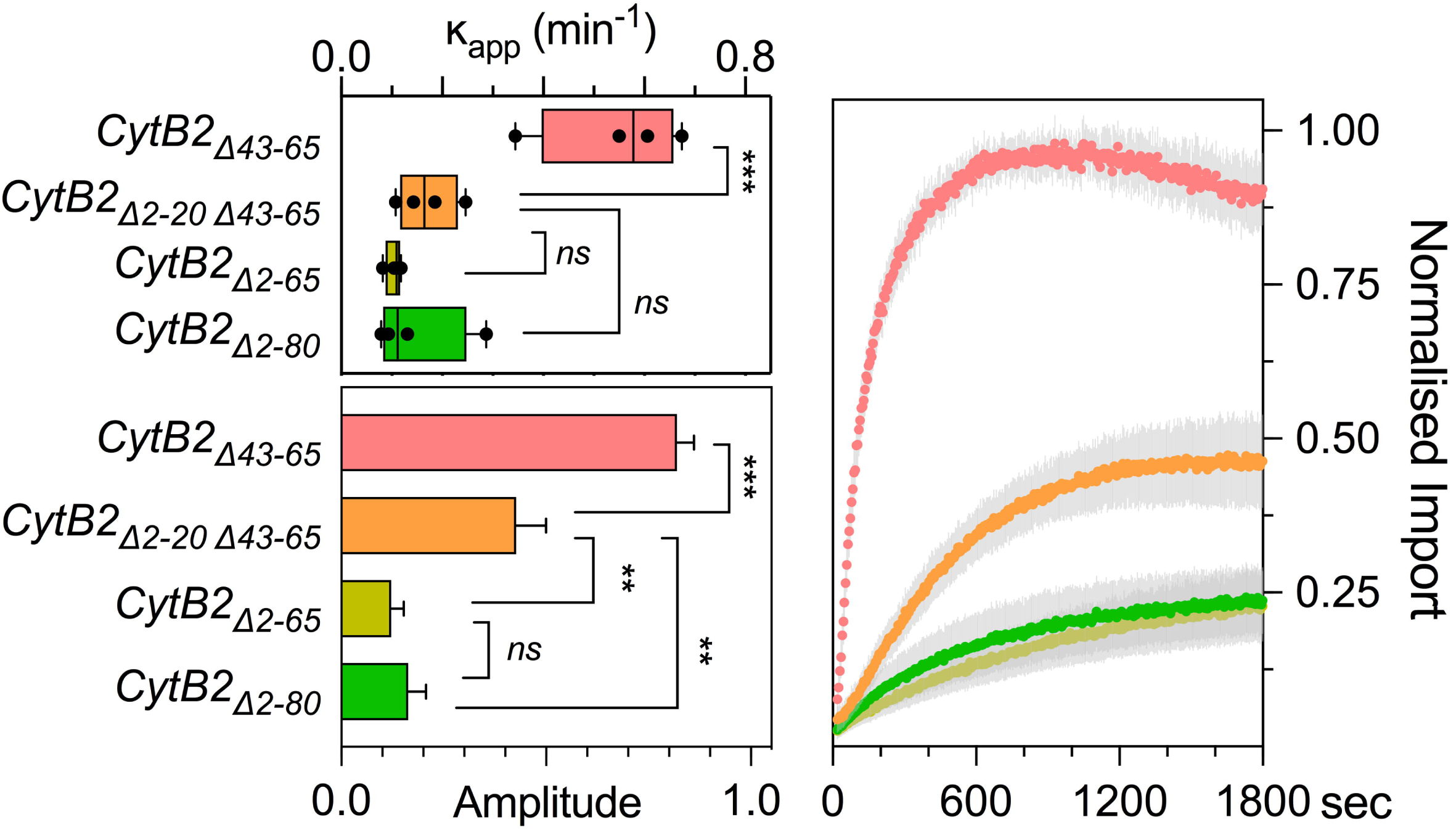
Effect of signal sequence deletion on mitochondrial import. The mitochondrial targeting sequence of CytB2_Δ43-65_-pep86 was truncated at different positions and import measured at 1 µM precursor in fully energised mitochondria (NADH + ATP regeneration system). Data represents 3-4 independent experiments and is shown as mean ± SEM. Differences between groups were analysed by a one-way ANOVA with predefined contrasts corrected with Holm-Sidak test. ***, p < 0.001; **, p< 0.01; ns, not significant.

## DISCUSSION

To understand protein transport across membranes, it is vital to be able to measure it. Until now, such experiments have been difficult to perform and scale, and have yielded very little kinetic insight. The splitluciferase-based assay optimised and validated here solves these problems. Indeed, our success in adapting NanoBiT to different conditions, both in bacteria and mitochondria, suggests that it will be compatible with many other membrane and non-membrane bound protein translocation machines.

Attempts to develop real-time import assays have been made previously, *e.g.* by labelling pre-proteins with fluorescent dyes^21,22^. These have provided some information on the mechanism of protein secretion by the bacterial Sec system, however, the risk of inefficient labelling represents possible competition for translocation by non-labelled protein, affecting overall kinetic analyses. And perhaps most importantly, fluorescent dyes are significantly different from amino acid side chains both in terms of size and composition – raising questions as to the physiological relevance of the measurements. The pep86 tag circumvents these problems: it can be cloned easily onto the C-terminus of a pre-protein, and as a short peptide it resembles exactly a native translocation substrate. Its relatively small footprint should not affect transport kinetics appreciably. Although we decided to place pep86 on the C-terminus of pre-proteins, the tag will bind with high affinity to 11S as long as it is available and not sterically hindered, which means it could also be placed in internal protein loops, as previously reported^23^.

Bipartite systems, such as the split-NanoLuc used in present work, have been used to evaluate protein translocation before^24^^-^^26^. Ensembled^27^ or split-GFP^25^ are limited by the fact that maturation of the chromophore (or that of any fluorescent protein) is a slow process, in the order of minutes^28^. This is much slower than protein translocation, and therefore would be rate limiting, precluding any kinetic assessment of the biological process, and probably explains why it has been only really used for low time- and spatial-resolution *in vivo* analysis of protein localisation. Split-β-galactosidase systems, such as CAPT^24^ or PathHunter^26^ (trademark of DiscoverX) are closer to our split-NanoLuc-based assay in that both rely on inactive fragments to restore enzyme activity. However, while Wehrman, et al. ^24^ reported a tag of 46 aa in the CAPT system that spontaneously associates with a bigger fragment, the authors did not measure binding affinity, so the response time may be limited by the association of the fragments.

A possible disadvantage when using high 11S concentrations is fast depletion of its substrate Furimazine. However, we lowered the amount of biological sample to avoid this problem while maintaining high sensitivity. It also suggests that in special cases where the amount of starting material for cell fractioning is an issue, such as mammalian cells, zebra fish, fruit flies or flat worms, our assay can be a practical tool to use.

The real-time assay showed a robust, reliable and high-dynamic range whenever it was challenged with standard controls for studying protein translocation. The signal readout, *i.e.* translocation was energy-dependent and signal-sequence specific and, it could be unambiguously inhibited with known inhibitors of the translocon. An important aspect of the system is its ability to distinguish nuances beyond the capabilities of the classical methods; either in terms of signal sequences or, to detect protein import in response to (patho)physiological energisation conditions. For example, to our knowledge it is the first time that protein import driven by the reversal of ATP synthase has been observed. In a cellular context, it means that respiration-impaired mitochondria can still efficiently import proteins as long as cytosolic ATP is available.

Contrarily to some recently developed high sensitivity methods^29-32^ for measuring translocation (Table S1), the requirements for our new assays is a simple luminometer, and standard molecular cloning procedures. Overall, this means that the real-time assay can be easily adopted by current or new laboratories working on protein translocation without the need of advanced knowledge on specific tools or techniques, while still providing high sensitivity and specificity.

The flexibility of the new assay show that it is not restricted to the systems presented here, but can be readily adapted to other frameworks, such as peroxisomes, chloroplasts, endoplasmic reticulum, nucleus or plasma membrane. Or even for monitoring protein translocation in non-membrane associated systems, such as to the interior of protein cages like GroEL or the proteasome.

Monitoring protein secretion through the plasma membrane using NanoBiT has already been described in mammalian cells^33^ and in Gram-positive bacteria^34^. However, employing the same approach to Gram-negative bacteria is considerably challenging because the outer membrane in their cell envelope provides a low permeability barrier. Therefore, instead of exogenous addition of 11S we targeted it to the periplasm while co-expressing the ‘*dark*’ peptide in the cytosol. This approach is preferable to the use of low affinity tags/peptides^9^^,^^24^ because it ensures that spontaneous binding in the destination compartment is retained and paves the way to monitor β-lactamases secretion *in vivo* and in real-time. Thus, we believe the assay will prove to be valuable for both fundamental research in protein trafficking systems, as well as for high throughput drug screening.

## MATERIAL AND METHODS

### Reagents

All chemicals, such as antibiotics, inducers and mitochondrial poisons, were of the highest commercially available grade of purity and were purchased from Sigma-Aldrich. Aqueous solutions were prepared in ultrapure (type I) water (Milli-Q Biocel A10 with pre-treatment via Elix 5, Millipore, Billerica, MA, USA). For non-aqueous solutions, ethanol (99.5%) or dimetylsulfoxide (DMSO), both from Sigma-Aldrich, were used as solvent.

### Culture conditions

Bacterial strains were cultured in LB or 2XYT for *in vivo* and *in vitro* experiments, respectively, with appropriate antibiotics (100 µg/ml ampicillin, 34 µg/ml chloramphenicol, 50 µg/ml kanamycin). Standard culturing temperature for *E. coli* MM52 (contains a temperature-sensitive genomic copy of SecA) was 30°C and 37°C was used for all other strains. Competent cells were prepared and transformed by heat-shock through standard procedures (Chung, Niemela, & Miller, 1989), with a 30 s incubation at 42°C and recovery in LB only.

Wild-type yeast, *Saccharomyces cerevisiae* strain YPH499, were cultured in standard YPD (1% yeast extract, 2% peptone, 2% glucose) at 30°C. Yeast mutants were cultured in synthetic complete growth media lacking uracil and supplemented with 2% glucose at 30°C. For liquid cultures, media was further supplemented with 0.0025% penicillin and 0.0025% streptomycin. Competent cells were prepared as previously described ^35^ and transformed by the LiAc/PEG method. Briefly, competent cells were rinsed in water and incubated in transformation mix containing 0.5 µg DNA, 3 µg/mL salmon sperm DNA (Sigma-Aldrich, UK), 100 mM LiAc, 10 mM Tris pH 7.5, 1 mM EDTA and 40% PEG 3000, for 30 min at 30°C followed by 15min at 42°C. Cells were let to recover in YPD for 60 min at 30°C before plating on synthetic growth media lacking uracil (Kaiser mixture; Formedium, UK) and supplemented with 2% glucose.

### Cloning

#### mt-11S for mitochondrial preparations

To produce yeast expressing 11S in the mitochondrial matrix, the 11S amino acid sequence previously published ^9^ was codon-optimised for *S. cerevisiae* and supplemented with the mitochondrial signal sequence of yeast alpha subunit of ATP synthase (ATP1/YBL099W; 1-35aa) on its N-terminus. This gene was purchased on a plasmid from Eurofins (Germany), digested with HindIII and XbaI, and ligated into the corresponding sites of pYES2CT (yielding pYES—mt-11S). The plasmid was verified by sequencing then transformed into YPH499 yeast cells. The *mt-11S* gene was cloned into a high-copy number plasmid (pYES2) under the control of GAL promoter to facilitate the delivery of high quantities of 11S to the mitochondrial matrix.

#### His-tagged 11S for proteoliposomes preparations

To produce and purify 11S, the *11S* gene (without the mitochondrial signal sequence) was amplified from pYES—mt-11S with a 5′ primer containing an NcoI site followed by a 6-his tag (GATCGTCCATGGGCCATCATCATCATCATCATGGCGTTTTCACATTGGAG), and a 3′ primer containing a HindIII restriction site (GCCTAAAAGCTTCTAGCTATTGATGGTTACACG). The resulting PCR product was digested with NcoI and HindIII, and ligated into the corresponding sites of pBAD/*Myc*-His C (yielding pBAD—_6H_11S). The plasmid was verified by sequencing then transformed into BL21(DE3) cells.

#### Tethered 11S for IMV preparations

To produce IMVs with high concentrations of 11S on the inside, we tethered 11S to the periplasmic face of the inner membrane using a lipid anchor^12^. The 11S gene (without the his-tag) was amplified from pBAD— _6H_11S using a 5′ primer with a NcoI restriction site followed by the signal sequence and first six amino acids of NlpA (EG10657; ACGTAGCCATGGGCAAACTGACAACACATCATCTACGGACAGGGGCCGCATTATTGCTGGCCGGAATTCTGCTGGCAGGTTGCGACCAGAGTAGCAGCGGCGTTTTCACATTGGAG), and a 3′ primer including the HindIII restriction site (CTACGTAAGCTTCTAGCT). The PCR product was then digested with NcoI and HindIII, and ligated into the corresponding sites in pRSFDuet-1. The resulting plasmid was verified by sequencing then cotransformed with pBAD—SecYEG into BL21(DE3) cells.

#### GST-dark peptide for *in-vitro* experiments

To allow recombinant expression and purification of large quantities of dark peptide, we fused it to the C-terminus of glutathione-S-transferase (GST). This was done by PCR insertion of a DNA sequence coding for a pep86 version that does not luminesce (dark peptide, see details in ^13^) immediately after the BamHI site in pGEX-1 (primer sequences: CATCCTCCAAAATCGGATCCCGGAGTGAGCGGCTGGGCGCTGTTTAAAAAAATTAGCTAAGAATTCATCGTGACTGAC and GTCAGTCACGATGAATTCTTAGCTAATTTTTTTAAACAGCGCCCAGCCGCTCACTCCGGGATCCGATTTTGGAGGATG). The resulting plasmid (pGEX—GST-dark) was verified by DNA sequencing, then transformed into BL21 (DE3) cells.

#### proOmpA(±pep86)

proOmpA with a C-terminal minimal V5 epitope was produced as described previously ^2^. For the real-time translocation assays, PCR insertion was used to add the pep86 sequence, preceded by a short GSG linker, after the V5 tag (primers: GAATCCGCTGCTGGGCCTGGGCTCCGGCGTGAGCGGCTGGCGCCTGTTTAAAAAAATTAGCTAAGCTTACGTAGAACAAAAAC and GTTTTTGTTCTACGTAAGCTTAGCTAATTTTTTTAAACAGGCGCCAGCCGCTCACGCCGGAGCCCAGGCCCAGCAGCGGATTC). After verifying the clone by DNA sequencing, proOmpA-pep86 was expressed and purified using exactly the same protocol as standard proOmpA.

#### proSpy(±pep86) and mSpy(±pep86)

To produce the periplasmic chaperone proSpy (EG13490), an *E. coli* optimised gene string (Life Technologies) for proSpy with a C-terminal minimal V5 epitope, a linker with pep86 tag, a TEV cleavage sequence and a 6-his tag was cloned into pBAD-HisA using overlap extension mutagenesis. For the mature form of Spy, a separate gene string was purchased omitting the signal sequence (1-23 aa). The resulting plasmids, pBAD —proSpy-pep86 and pBAD—mSpy-pep86, were verified by DNA sequencing, then transformed into MM52 and BL21(DE3), respectively.

#### CytB2±pep86 and Δmts-CytB2-pep86

An engineered version of yeast cytochrome B2 (YML054C, ^18^) comprising the first 158 aa with its hydrophobic domain on the signal sequence deleted (Δ43-65) followed by two tandem TEV cleave sites, a Myc tag and a C-term 6xhis tag, was codon-optimised for *E. coli* and purchased as a gene on plasmid (Eurofins, Germany). Then, the plasmid was digested with NcoI and HindIII, and the insert ligated into the corresponding sites of pBAD/*Myc*-His C (yielding pBAD--CytB2_Δ43-65_). For the real-time translocation assays, PCR insertion was used to add the pep86 sequence, preceded by a SGGGGS linker, after the 6xhis tag, yielding pBAD--CytB2_Δ43-65_-pep86_._ All plasmids were verified by sequencing then transformed into BL21(DE3) cells.

For CytB2 with different truncations on its mitochondrial targeting sequence, an *E. coli* optimised sequence of CytB2 comprising the first 220 aa and the Δ43-65 deletion was supplemented with a Myc tag, a 6xhis tag followed by a GGGS linker and a C-term pep86 tag. The sequence was purchased as gene on plasmid (Eurofins) and digested with NcoI and HindIII. Then, the insert was ligated into the corresponding sites of pBAD/*Myc*-His C (yielding pBAD--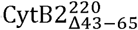 – pep86). The different MTS truncations were achieved by plasmid PCR deletion using pBAD--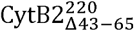 – pep86 as template and the following primers to yield the corresponding plasmids: GCGAGCAAAACCCGGTTAAATAC and CATGGTTAATTCCTCCTGTTAGCC for pBAD-- 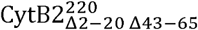 – pep86; TCTAGTGTTGCGTATCTGAATTGG and CATGGTTAATTCCTCCTGTTAGCC for pBAD-- 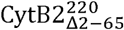 – pep86, and; GAACCGAAACTCGACATGAACAAA and CATGGTTAATTCCTCCTGTTAGCC for pBAD--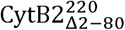 – pep86. All plasmids were verified by sequencing then transformed into BL21(DE3) cells.

#### pro11S--GST-dark for *in-vivo* experiments

To produce bacteria expressing 11S in the periplasm, the amino acid sequence of 11S was codon-optimised for *E. coli*, supplemented with the signal sequence of bacteria OmpA (EG10669, 1-21 aa) on its N-terminus (pro 11S) and purchased as a gene fragment (GeneArt, Invitrogen). The fragment was ligated into pMiniT2.0 according to the manufacturer’s instructions (NEB PCR cloning kit, E1203S), and sequence checked with pMiniT F and R primers. pMini—pro11S and pBAD-Myc-His-C were digested with NcoI and HindIII (NEB HF enzymes) according to the manual, fragments agarose gel purified (QIAquick gel extraction kit, 28704), ligated using T4 ligase (ThermoScientific, #EL0011) and transformed into α-select creating pBAD—pro11S. The GST-dark gene fragment was synthesised (Invitrogen), ligated into pMiniT2.0 according to the manual and sequence checked. pMini—GST-dark and pBAD—pro11S were digested with HindIII and SalI (NEB HF enzymes) according to the manual, fragments agarose gel purified, ligated using T4 ligase and transformed into *E. coli* α-select creating pBAD—pro 11S—GST-dark.

#### NDM1-pep86 for *in-vivo* experiments

The pep86 tag was added to the 3’ end of *ndm*-1 by site-directed mutagenesis PCR (Phusion Site-Directed Mutagenesis Kit, ThermoScientific, #F541) using 5’ phosphorylated primers (forward: 5’ PHO-CTGTTTAAAAAAATTAGCTAAGCCATGGCTGACCACGTCACC and reverse: 5’ PHO-GCGCCAGCCGCTCACGCCGCTGCCGCGCAGCTTGTCGGC) according to manual (64°C annealing temperature for 30s, 90s extension) and pSU2718-NDM1 as template. The circularised plasmid was ligated according to manufacturer’s instructions and transformed into *E. coli* α-select creating pSU2718-NDM1-pep86.

### Protein expression and purification

For expression, pre-cultures were inoculated with a single colony of the desired bacterial strain (BL21(DE3) as default) and grown in LB with appropriate antibiotic for 16 h at 37°C, 200 rpm. Cultures were inoculated at 1:100 from pre-cultures in 2xYT plus antibiotic and grown at 37°C, 200 rpm until mid-log phase, then induced for 2½-3 h with 0.1-0.2% (w/v) arabinose or 1 mM depending on the plasmid. For overexpression of bacterial pre-proteins the MM52 strain was used instead as it contains a temperature-sensitive copy of genomic SecA^16^. When exposed to temperatures above 30°C the mutant SecA is rendered inactive, causing pre-proteins to accumulate in the cytoplasm, typically as inclusion bodies. Therefore, pre-cultures were grown at 30°C and protein expression carried out at 39°C.

#### His-tagged 11S

Cells were harvested, resuspended in 20 mM Tris pH 7.5, 50 mM KCl (TK) with 10% glycerol (TKG) then cracked open using a cell disruptor (Constant Systems) and clarified by centrifugation. The supernatant was loaded onto a Ni^2+^ column packed with chelating Sepharose Fast Flow resin (GE Healthcare), washed in TKG with 50 mM imidazole, then eluted with TKG + 330 mM imidazole. Imidazole was removed by washing with TKG in a spin concentrator, and the final protein concentration determined from A280, using the calculated extinction coefficient of 19,940 M^-1^.cm^-1^. The sample was then snap frozen and stored at -80°C.

#### GST-dark peptide

Cells were harvested, resuspended in TK buffer then cracked in a cell disruptor and clarified by centrifugation. The supernatant was loaded onto a GSTrap 4B (GE Healthcare) at 4 °C and the column washed with TK buffer until A_280_ of the flowthrough stopped decreasing. Elution was performed with 10 µM reduced glutathione in TK. The yield of the resulting protein (hereafter GST-dark) was determined from A_280,_ using the calculated molar extinction coefficient of 48,360 M^-1^.cm^-1^. The sample was then snap frozen and stored at -80°C.

#### pep86-tagged mitochondrial precursors

Cells were harvested, resuspended in TK buffer then cracked in a cell disruptor and clarified by centrifugation. Inclusion bodies were solubilised in TK plus 6 M urea (TK+urea) before loading into an in-house packed Ni^2+^ column. After washing with TK+urea, proteins were eluted with 330 mM imidazole in TK+urea and loaded into an in-house packed Q- or S-column. After column wash, proteins were thereafter eluted in TK+urea+1 M KCl gradient (0-100%) during 20 min. Final fraction was spin concentrated and the final protein concentration determined from A_280,_ using the calculated extinction coefficient. The sample was then snap frozen and stored at -80°C.

#### Pep86-tagged bacterial pre-proteins

For **proOmpA**, the cell pellet was resuspended in 130 mM NaCl, 20 mM Tris pH 8.0 and cells cracked in a cell disruptor followed by a clarifying spin. A previously established purification protocol was utilised where inclusion bodies were harvested by gentle centrifugation at 4000 g for 15 minutes and solubilised in urea^2^. The resulting mixture was loaded onto an anion exchange column equilibrated in a salt-free 6 M urea, 10 mM Tris pH 8.0 buffer. A linear salt gradient of 0-1 M was then applied, where proOmpA-pep86 constituted the first protein to elute, with a 280 nm absorbance peak at approximately 40 mM NaCl.

For **proSpy**, cells were harvested and resuspended in lysis buffer (500 mM NaCl, 50 mM Tris, 30 mM imidazole, pH 8.0) buffer supplemented with cOmplete, EDTA-free Protease Inhibitor Cocktail. The cells were then lysed and clarified by centrifugation. The soluble cell fraction was determined to contain approximately 80% of the total expressed Spy and was therefore loaded onto a 5 mL HiTrap Crude FF nickel affinity chromatography column (GE Healthcare). After washing with lysis buffer, the bound proteins were eluted with lysis buffer containing 300 mM imidazole and then spin-concentrated (5 KDa cut-off) to ~14 mL before TEV digestion. DTT and EDTA, 1 mM and 0.5 mM respectively, plus ~0.1 mg/mL TEV protease were added to the suspension and incubated at room temperature for about 3 h. Then, Spy solution was purified by nickel affinity chromatography as described above, but this time collecting the unbound column flow through (His tag removed-Spy). The sample was dialysed overnight into 6 M urea, 20 mM Tris pH 8.0 and then snap frozen and stored at -80°C.

### Proteoliposome (PL) preparation

SecYEG proteoliposomes were produced as described previously ^3^. Briefly, purified SecYEG was mixed with *E. coli* polar lipids in DDM, then the detergent removed gradually using BioBeads. To encapsulate 11S, we simply included purified 11S at the desired final concentration (20 µM standard, or as noted in the text) in the SecYEG/polar lipid mix, prior to the addition of biobeads. For the initial experiments, SecYEG/11S PLs were harvested by centrifugation (30 mins at 100,000 g) as per the standard method, washed twice by resuspending in 3 mL TKM then centrifuging for 30 min at 100,000 g and pipetting off the supernatant, then resuspended to give the desired final SecYEG concentration. These additional washing steps removed most of the non-encapsulated 11S, reducing the background signal for the transport assays – although they did not obviate the need for GST-dark. For the 11S concentration series we instead passed the reconstituted PLs over a gravity flow Sephacryl-S1000 column to separate away unbound 11S. These PLs were quantified by scattering at 320 nm (relative to PLs produced using the standard method), then used directly in transport assays. This method is both more effective at removing free 11S and eliminates the centrifugation and resuspension steps, which potentially damage PLs and cause them to leak.

### IMV preparation

IMVs were either prepared from *E. coli* BL21 (DE3) or a strain lacking ATP synthase (unc^-^; HB1 cells^36^), to prevent PMF generation upon addition of ATP. Cells were grown to mid-log phase and 37 °C in 2xYT supplemented with 100 µg/mL ampicillin and 50 µg/L kanamycin, then co-induced for 2½ h with 0.1% arabinose and 1 mM IPTG. Inverted membrane vesicles were prepared from the membranes as described previously ^2^.

### Mitochondrial isolation

mt-11S-expressing yeast were grown overnight at 30°C in synthetic growth media lacking uracil and supplemented with 3% glycerol plus 0.0025% Pen/Strep. Yeast cells were cultured in glycerol-based media to increase mitochondrial mass and maximise mitochondrial function ^19^. 1% galactose was added at mid-log phase to start inducing mt-11S (total time ~16h). In the end, mitochondria were isolated through differential centrifugation after cell wall was reduced by 1 mM DTT in 100 mM Tris-SO_4_ buffer for 15 min at 30°C and then digested with zymolase in sorbitol-phosphate buffer for 30 min at 30°C. The final mitochondrial pellet was resuspended in 250 mM sucrose and 10 mM MOPS, pH 7.2. Mitochondrial protein was quantified by BCA assay, using BSA as standard.

### Binding experiments

Complementation of pep86-tagged precursor with pure _6H_11S was performed in 1x Nano-Glo buffer diluted with TK buffer and Prionex (0.1% final). The reaction mix containing 1x furimazine was used to prepare a titration curve of precursor ranging from 13.9 nM to 3 µM which was added to a 96-well plate. Separately, pure 11S was diluted in reaction mix supplemented with 1x furimazine so that automatic injection of 80 µL would give the desired concentration upon addition (30 pM in 100 µL). Luminescence was read on BioTek Synergy Neo2 plate reader (BioTek Instruments, UK) without emission filters every 0.5 sec during 30 sec at 25°C working on well mode. Obtained data was fitted to a single exponential.

### Western blot transport assays

For the **Sec-system**, western blot transport time courses were performed in a 25 °C heat block. Reaction master mixes were prepared in buffer TK + 2 mM MgCl_2_ (TKM), containing: creatine phosphate to 5 mM, creatine kinase to 0.1 mg/mL, SecYEG IMVs or PLs to 4% of final volume, SecA to 1 µM and proOmpA to 0.2 µM. From each master mix, 5 µL was set aside for a 10 % control, and another 50 µl diluted 5-fold into ice-cold 1 mg/mL protease K in 5 mM EDTA as a-ATP (t=0) control. Reactions were immediately started by adding ATP to a final concentration of 1 mM, and 50 µL was taken at various time points and quenched rapidly as for the t=0 sample.

Samples were prepared and western blotted essentially as described previously^2^. Briefly, undigested protein was precipitated on ice for 30 mins with 20% (w/v final) trichloroacetic acid, then centrifuged and the supernatant removed. Pellets were dried in a vacuum centrifuge, then resuspended in 30 µL 2.5x NuPage LDS buffer (Thermo Fisher Scientific) and heated to 70 °C for 20 mins. 10 µL of the resulting samples were run out on a gel, blotted, then developed using an α-v5 primary antibody (SV5-Pk1, GeneTex) and a DyLight 800-labelled secondary antibody (Thermo Fisher Scientific). Western blots were visualised on an Odyssey Fc (LI-COR) and the bands quantified using the built-in software.

For the **mitochondrial system**, yeast mitochondria (240 μg protein) were diluted in 240 μL import buffer (250 mM sucrose, 80 mM KCl, 5 mM MgCl_2_, 10mM K_2_HPO_4_, 10 mM MOPS-KOH pH 7.2) supplemented with 2 mM NADH, 2 mM ATP, 5 mM creatine phosphate and 0.1 mg/mL creatine kinase. The samples were preincubated at 25 °C for 5 min before import was started by adding 1 μg/mL substrate. The reaction was allowed to proceed for 1, 3 or 5 min with gentle shaking at 350 rpm before being stopped by the addition of 2.4 uL VOA (containing 100 μM valinomycin, 2 mM Oligomycin and 800 μM antimycin A). Half of each sample was treated with 25 μg/mL Proteinase K on ice for 15 min followed by the addition of 3 mM phenylmethylsulfonyl fluoride (PMSF) for 2-5 min to stop the reaction. Centrifugation at 20,000 g was used to isolate the mitochondria which were then washed with SM buffer (250 mM sucrose, 10 mM MOPS-KOH, pH7.2). Pellets were resolved in 2x sample buffer (4% SDS, 20% glycerol, 125 mM Tris-HCl pH 6.8, 0.02% bromophenol blue, 50 mM DTT) and heated at 65 °C for 10 min before being subjected to SDS-PAGE. After blotting, membranes were developed using an anti-myc primary antibody (Cell Signaling) and a DyLight 800-labelled secondary antibody (Thermo Fisher Scientific). Western blots were visualised on an Odyssey Fc (LICOR) and the bands quantified using the built-in software.

### Real-time import assay

#### Cuvette mode

Real-time import assays (for the Sec system) were performed at 25 °C in a Jobin Yvon Fluorolog (Horiba) with the lamp turned off and emission measured at 460 nm (with slits open to maximum, i.e. 10 nm bandpass). A reaction mix was assembled in a 1 mL cuvette with a stirrer bar by adding (in order): TKM to give a final volume of 1 mL, Prionex (Sigma-Aldrich; registered trademark of Pentapharm AG, Basel) to 0.1%, 10 µL Nano-Glo substrate (furimazine, Promega), creatine phosphate to 5 mM, creatine kinase to 0.1 mg/mL, GST-dark to 40 µM, 1 µL SecYEG/11S HB1(DE3) IMVs or PLs, and SecA to 1 µM. After a 5 min equilibration, a luminescence baseline signal was measured for 1 minute, followed by addition of proOmpA-pep86 to 1 µM final concentration. After a further 10 minutes, ATP was added to 1 mM final concentration, and the transport reaction followed until completion.

The background, caused by association of proOmpA-pep86 with non-encapsulated 11S, fits well to a single (PLs) or double (IMVs) exponential (Fig. 2). Therefore, we fitted the signal from after proOmpA-pep86 but prior to ATP addition to a single exponential, and subtracted the resulting fit from the raw data. This corrected data corresponds to ATP-driven protein transport.

#### Microplate mode

For the **Sec-system**, all reactions were carried in 20 mM Tris, 50 mM KCl, 2 mM MgCl2, pH 8.0 (TKM)supplemented with 0.1% Prionex, 0.1 mg/mL creatine kinase, 5 mM creatine phosphate, 40 µM GST-dark, 1x Nano-Glo substrate (furimazine, Promega), 0.3 µM SecA and at 25°C in a low-binding white 96 well plate. Reaction wells were setup with 3 µL of pre-protein (final concentration approximately 3 µM) and, immediately prior to starting measurements, 97 µL of master mix, such that on automated injection of 25 µL of 5 mM ATP solution the above concentrations were achieved. Background luminescence (likely caused by 11S from damaged IMVs) was measured for 10 min on a BioTek Synergy Neo2 plate reader (BioTek Instruments, UK) until equilibration was achieved (steady-state luminescence), at which point 25 µL of ATP was automatically injected and mixed thoroughly (for 2 seconds) to initiate translocation. Luminescence was collected without emission filters for an additional 20 min, with the lowest interval time possible for the number of samples being measured.

For the **mitochondrial system**, reactions were carried out in 300 mM mannitol, 10 mM HEPES pH 7.4, 25 µM EGTA, 1 mM KH_2_PO_4_ supplemented with 0.1% Prionex, 10 µM GST-dark, 2 mM NADH, 25-50 µg/mL of frozen yeast mitochondria and 0.25x Nano-Glo substrate (furimazine, Promega), at 25°C in a low-binding white 96 well plate. Creatine kinase (0.1 mg/mL), creatine phosphate (5 mM) and ATP (1 mM) were also included in the buffer unless stated otherwise. Reactions started by the addition of 25 µL of precursor to make a final volume of 125 µL. When plates were read on a Packard Lumicount BL10001 (Packard BioSciences, Meriden, CT, US), additions were made manually using an 8-channel pipette with manual mixing; if BioTek Synergy Neo2 plate reader (BioTek Instruments, UK) was used instead, additions were made automatically using the system pump set to the default injection speed followed by a 5 sec linear shaking step. In both plate readers, luminescence was collected for 0.2 sec/well, without emission filters, and the gain was set to allow maximum sensitivity without detector saturation. An initial baseline of 60 sec was acquired before precursor addition and then luminescence was read for at least 20 min. Time between reads was set to the minimum allowed interval in each plate reader – 10-12 sec for LumiCount and 5 sec for Neo2.

### *In-vivo* β-lactamase secretion assay

*E. coli* MC4100 were transformed with pBAD—pro11S--GST-dark alone or in combination with pSU2718--NDM-1-pep86. Starter cultures were inoculated with a single colony of the desired strain and grown in LB with appropriate antibiotic for 16h at 30°C, 200 rpm. 5 mL cultures (LB with antibiotic) were inoculated from these starter cultures at 1:100 and grown at 30°C, 200 RPM until OD_600_ ~ 1. At that point, all cultures were induced by adding arabinose to a final concentration of 0.2% (w/v) and incubated at 39°C. 100 µL of sample was taken at the time of induction and assayed 2 h later using the Nano-Glo Live Cell Assay System (Promega) according to the manufacturer’s protocol. Luminescence was measured over 100 reads during 10-20 min and the data averaged. Background luminescence was deducted from assay data of *bla* carrying strains.

### Data analysis and Statistics

Results are shown as means ± SEM of the indicated number of experiments. Apparent rates (*k*_app_) were calculated as the reciprocal of the time it takes to reach half of the maximal luminescent signal (t_50%_). Statistical significance between mean differences was determined using two-tailed Student’s t test or one-way ANOVA, when more than two groups were analysed, followed by predefined contrasts using Bonferoni’s post-hoc analysis to correct for multiple comparisons. Differences were considered significant if p < 0.05 and categorized accordingly to their interval of confidence. Statistical analyses were performed using Graph Pad Prism version 8.0.0 (GraphPad Software, Inc., San Diego, CA, USA).

## ACKNOWLEDGMENTS

This work was primarily funded by Wellcome (Investigator Award for GCP, WJA, DN, XL, AR and IC; 104632) and the BBSRC: BB/N015126/1 (project grant, DWW and IC) and BB/L01386X/1 (BrisSynBio, LB and IC). We thank Prof. M. Avison for the gift of the NDM1 expression plasmid.

## AUTHOR CONTRIBUTIONS

GCP designed, executed and analysed data for the real-time assay in mitochondria. WJA and DWW designed, executed and analysed data for the bacterial Sec system. LB and IC performed and designed the in vivo experiments, respectively. XL and DN were responsible for protein purification. AR performed and analysed experiments for the mitochondrial system using conventional methods. GCP, WJA, DWW, AC and IC wrote the manuscript.

## Abbreviations

AA: antimycin A
IMM: inner mitochondrial membrane
mt-11S: mitochondrial-targeted 11S
_H6_11S: N-terminal 6xHis tagged version of 11S
OMM: outer mitochondrial membrane
OVA: oligomycin, valinomycin and antimycin A (full depolarisation cocktail)
MTS: mitochondrial targeting sequence
NanoBiT: NanoLuc Binary Technology
PAM: presequence-translocase-associated import-motor
pep86: high-affinity version of NanoBiT small fragment
pep114: small fragment of NanoBiT
PMF: proton motive force
TIM: translocase of the inner membrane
TOM: translocase of the outer membrane
11S: large fragment of the NanoBiT
ΔΨ: membrane potential.

## REFERENCES

1 Song, C., Kumar, A. & Saleh, M. Bioinformatic comparison of bacterial secretomes. Genomics Proteomics Bioinformatics 7, 37-46, doi:10.1016/S1672-0229(08)60031-5 (2009).

2 Corey, R. A. et al. Specific cardiolipin-SecY interactions are required for proton-motive force stimulation of protein secretion. Proc Natl Acad Sci U S A 115, 7967-7972, doi:10.1073/pnas.1721536115 (2018).

3 Gold, V. A., Robson, A., Clarke, A. R. & Collinson, I. Allosteric regulation of SecA: magnesium-mediated control of conformation and activity. J Biol Chem 282, 17424-17432, doi:10.1074/jbc.M702066200 (2007).

4 Miyata, N. et al. Pharmacologic rescue of an enzyme-trafficking defect in primary hyperoxaluria 1. Proc Natl Acad Sci U S A 111, 14406-14411, doi:10.1073/pnas. 1408401111 (2014).

5 Miyata, N. et al. Adaptation of a Genetic Screen Reveals an Inhibitor for Mitochondrial Protein Import Component Tim44. J Biol Chem 292, 5429-5442, doi:10.1074/jbc.M116.770131 (2017).

6 Brundage, L., Hendrick, J. P., Schiebel, E., Driessen, A. J. & Wickner, W. The purified E. coli integral membrane protein SecY/E is sufficient for reconstitution of SecA-dependent precursor protein translocation. Cell 62, 649-657 (1990).

7 Gorlich, D. & Rapoport, T. A. Protein translocation into proteoliposomes reconstituted from purified components of the endoplasmic reticulum membrane. Cell 75, 615-630 (1993).

8 Kang, P. J. et al. Requirement for hsp70 in the mitochondrial matrix for translocation and folding of precursor proteins. Nature 348, 137-143, doi:10.1038/348137a0 (1990).

9 Dixon, A. S. et al. NanoLuc Complementation Reporter Optimized for Accurate Measurement of Protein Interactions in Cells. ACS Chem Biol 11, 400-408, doi:10.1021/acschembio.5b00753 (2016).

10 Lill, R. et al. SecA protein hydrolyzes ATP and is an essential component of the protein translocation ATPase of Escherichia coli. EMBO J 8, 961-966 (1989).

11 Tani, K., Shiozuka, K., Tokuda, H. & Mizushima, S. In vitro analysis of the process of translocation of OmpA across the Escherichia coli cytoplasmic membrane. A translocation intermediate accumulates transiently in the absence of the proton motive force. J Biol Chem 264, 18582-18588 (1989).

12 Harvey, B. R. et al. Anchored periplasmic expression, a versatile technology for the isolation of high-affinity antibodies from Escherichia coli-expressed libraries. Proc Natl Acad Sci U S A 101, 9193-9198, doi:10.1073/pnas.0400187101 (2004).

13 Dixon, A. S. V., WI, US), Encell, Lance (Fitchburg, WI, US), Hall, Mary (Waunakee, WI, US), Wood, Keith (Mt. Horeb, WI, US), Wood, Monika (Mt. Horeb, WI, US), Schwinn, Marie (Madison, WI, US), Binkowski, Brock F. (Sauk City, WI, US), Zegzouti, Hicham (Madison, WI, US), Nath, Nidhi (Madison, WI, US), Mondal, Subhanjan (Middleton, WI, US), Goueli, Said (Fitchburg, WI, US), Meisenheimer, Poncho (San Luis Obispo, CA, US), Kirkland, Thomas (Atascadero, CA, US), Unch, James (Arroyo Grande, CA, US), Pulukkunat, Dileep K. (Middleton, WI, US), Robers, Matthew (Madison, WI, US), Dart, Melanie (Madison, WI, US), Machleidt, Thomas (Madison, WI, US). ACTIVATION OF BIOLUMINESCENCE BY STRUCTURAL COMPLEMENTATION. United States patent (2014).

14 Eichler, J., Rinard, K. & Wickner, W. Endogenous SecA catalyzes preprotein translocation at SecYEG. J Biol Chem 273, 21675-21681 (1998).

15 Ckakraborty, A. et al. Molecular characterization and clinical significance of New Delhi metallo-beta-lactamases-1 producing Escherichia coli recovered from a South Indian tertiary care hospital. Indian J Pathol Microbiol 58, 323-327, doi:10.4103/0377-4929.162864 (2015).

16 Oliver, D. B. & Beckwith, J. E. coli mutant pleiotropically defective in the export of secreted proteins. Cell 25, 765-772 (1981).

17 Webster, K. A. & Bronk, J. R. Ion movements during energy-linked mitochondrial structural changes. J Bioenerg Biomembr 10, 23-44 (1978).

18 Gold, V. A. et al. Visualizing active membrane protein complexes by electron cryotomography. Nat Commun 5, 4129, doi:10.1038/ncomms5129 (2014).

19 Backes, S. et al. Tom70 enhances mitochondrial preprotein import efficiency by binding to internal targeting sequences. J Cell Biol 217, 1369-1382, doi:10.1083/jcb.201708044 (2018).

20 Klaus, C., Guiard, B., Neupert, W. & Brunner, M. Determinants in the presequence of cytochrome b2 for import into mitochondria and for proteolytic processing. Eur J Biochem 236, 856-861 (1996).

21 De Keyzer, J., Van Der Does, C. & Driessen, A. J. Kinetic analysis of the translocation of fluorescent precursor proteins into Escherichia coli membrane vesicles. J Biol Chem 277, 46059-46065, doi:10.1074/jbc.M208449200 (2002).

22 Liang, F. C., Bageshwar, U. K. & Musser, S. M. Bacterial Sec protein transport is rate-limited by precursor length: a single turnover study. Mol Biol Cell 20, 4256-4266, doi:10.1091/mbc.E09-01-0075 (2009).

23 Dixon, A. S. V., WI, US), Encell, Lance P. (Fitchburg, WI, US), Machleidt, Thomas (Madison, WI, US), Schwinn, Marie (Madison, WI, US), Wood, Keith (Mt. Horeb, WI, US), Wood, Monika (Mt. Horeb, WI, US), Zimmerman, Kris (Madison, WI, US). Internal protein tags. United States patent (2018).

24 Wehrman, T. S., Casipit, C. L., Gewertz, N. M. & Blau, H. M. Enzymatic detection of protein translocation. Nat Methods 2, 521-527, doi:10.1038/nmeth771 (2005).

25 Smoyer, C. J. et al. Analysis of membrane proteins localizing to the inner nuclear envelope in living cells. J Cell Biol 215, 575-590, doi:10.1083/jcb.201607043 (2016).

26 Patel, A. et al. A combination of ultrahigh throughput PathHunter and cytokine secretion assays to identify glucocorticoid receptor agonists. Anal Biochem 385, 286-292, doi:10.1016/j.ab.2008.11.005 (2009).

27 Calmettes, G. & Weiss, J. N. A quantitative method to track protein translocation between intracellular compartments in real-time in live cells using weighted local variance image analysis. PLoS One 8, e81988, doi:10.1371/journal.pone.0081988 (2013).

28 Iizuka, R., Yamagishi-Shirasaki, M. & Funatsu, T. Kinetic study of de novo chromophore maturation of fluorescent proteins. Anal Biochem 414, 173-178, doi:10.1016/j.ab.2011.03.036 (2011).

29 Rhee, H. W. et al. Proteomic mapping of mitochondria in living cells via spatially restricted enzymatic tagging. Science 339, 1328-1331, doi:10.1126/science.1230593 (2013).

30 Hoogewijs, K. et al. Assessing the Delivery of Molecules to the Mitochondrial Matrix Using Click Chemistry. Chembiochem 17, 1312-1316, doi:10.1002/cbic.201600188 (2016).

31 Ozawa, T., Sako, Y., Sato, M., Kitamura, T. & Umezawa, Y. A genetic approach to identifying mitochondrial proteins. Nat Biotechnol 21, 287-293, doi:10.1038/nbt791 (2003).

32 Ozawa, T. et al. A minimal peptide sequence that targets fluorescent and functional proteins into the mitochondrial intermembrane space. ACS Chem Biol 2, 176-186, doi:10.1021/cb600492a (2007).

33 Rouault, A. A. J., Lee, A. A. & Sebag, J. A. Regions of MRAP2 required for the inhibition of orexin and prokineticin receptor signaling. Biochim Biophys Acta Mol Cell Res 1864, 2322-2329, doi:10.1016/j.bbamcr.2017.09.008 (2017).

34 Wang, C. Y., Patel, N., Wholey, W. Y. & Dawid, S. ABC transporter content diversity in Streptococcus pneumoniae impacts competence regulation and bacteriocin production. Proc Natl Acad Sci U S A 115, E5776-E5785, doi:10.1073/pnas.1804668115 (2018).

35 Gietz, R. D. & Schiestl, R. H. High-efficiency yeast transformation using the LiAc/SS carrier DNA/PEG method. Nat Protoc 2, 31-34, doi:10.1038/nprot.2007.13 (2007).

36 Ballhausen, B., Altendorf, K. & Deckers-Hebestreit, G. Constant c10 ring stoichiometry in the Escherichia coli ATP synthase analyzed by cross-linking. J Bacteriol 191, 2400-2404, doi:10.1128/JB.01390-08 (2009).

